# Experiments on osmotically driven flow in idealized elastic membranes

**DOI:** 10.1101/2022.06.16.496432

**Authors:** Mazen Nakad, Jean-Christophe Domec, Sanna Sevanto, Gabriel Katul

**Affiliations:** Department of Civil and Environmental Engineering, Duke University, Durham, NC, USA; Bordeaux Sciences Agro, UMR 1391 INRA-ISPA, France; Nicholas School of the Environment, Duke University, Durham, NC, USA; Earth and Environmental Sciences Division, Los Alamos National Laboratory, Los Alamos, NM, USA

**Keywords:** Dextran, elastic membranes, M*ü*nch mechanism, osmosis, phloem, Van’t Hoff relation

## Abstract

The phloem provides a pathway for products of photosynthesis to be transported to different parts of the plant for consumption or storage. The M*ü*nch pressure flow hypothesis (PFH) is considered the leading framework to mathematically represent this transport. It assumes that osmosis provides the necessary pressure differences to drive the fluid flow and sucrose within the phloem. Mathematical models utilizing the PFH approximate the phloem by a relatively rigid semi-permeable tube. However, the phloem consists of living cells that contract and expand in response to pressure fluctuations. The effect of membrane elasticity on osmotically driven sucrose front speed has rarely been considered and frames the scope here. Laboratory experiments were conducted to elucidate the elastic-to-plastic pressure-deformation relation in membranes and their effect on sucrose front speeds. It is demonstrated that membrane elasticity acts to retard the sucrose front speed. The retardation emerges because some of the osmotic pressure performs mechanical work to expand the membrane instead of pressurizing water. These results offer a novel perspective about the much discussed presence of sieve plates throughout the phloem acting as structural dampers.

## 1 Introduction

A variety of fluid transport phenomena in plants do not require pumping or externally imposed pressure difference to move solutes and fluids. Instead, they rely on osmotic pressure to move loaded solutes and water simultaneously. A prototypical example is sucrose transport in the phloem of plants formulated in the 1930s by Ernst Münch and is now labeled as the pressure-flow hypothesis (Münch 1930) or PFH. The PFH is routinely used to describe the transport of photosynthates from production sites (leaves) to areas of consumption or storage (growing tissues and roots). This mechanism has been used to explain aspects of plant mortality under extreme weather conditions Sevanto et al. (2014) and ecosystem-scale impacts on carbon and water cycling Nikinmaa et al. (2013). Many models for phloem transport have been formulated and discussed (Phillips & Dungan 1993, Thompson & Holbrook 2003, Jensen et al. 2009, Cabrita et al. 2013, Sevanto 2014, Nakad et al. 2022).

One of the simplifications used in these models is that phloem is presented as a long cylindrical and relatively rigid slender tube with semi-permeable walls that enable water but not sugar to be exchanged between the sieve tube and its surroundings. Since the phloem is composed of living cells with weakly lignified cell walls (Sevanto et al. 2018), the rigid tube assumption may not be appropriate. Tube wall rigidity can impact sucrose transport rates through a number of mechanisms that remain to be uncovered (Tateshima et al. 2010). One latent mechanism is the dilution effect. Membrane expansion decreases the osmotic potential (i.e. the driving mechanism) by volumetrically increasing water storage within the membrane. In case of a rigid tube (or weakly elastic), any inflow of water is balanced by an equivalent outflow of water (due to conservation of mass) leading to a negligible water storage within the membrane and thus no dilution effects. One mathematical model estimated the impact of tube wall rigidity by assuming that volume expansion linearly relates to hydrostatic pressure within the phloem Thompson & Holbrook (2003). The model calculations suggest that when using typical values for the phloem’s Young modulus of elasticity, phloem elasticity had minimal impact on transport timescale. However, this model might underestimate the phloem elasticity because the Young modulus values were based on measurements of the whole phloem tissue (i.e. sieve tubes surrounded by other cell types in full turgor).

The work here employs experiments on idealized semi-permeable elastic membranes to explore generic relations between solute front speeds and membrane elasticity. These experiments are only intended to replicate the interplay between the physical mechanisms linking solute movement in the phloem to osmotic pressures from a structural point of view. As earlier noted, the phloem is composed of many elongated, cylindrical living cells interconnected by sieve plates. These plates are thin with large perforations to reduce flow resistance (Savage et al. 2017) and allow large molecules to pass. They are modified plasmodesmata and are consequence of cell development (Lucas et al. 1993, Kalmbach & Helariutta 2019). However, current studies consider these sieve plates as the “inevitable evil’ in terms of hindering phloem flow because they increase flow resistance (Savage et al. 2017, Jensen et al. 2012, Jensen 2018). Others (Lang 1979) attach a hydraulic function that supports flow because increased resistance may enable buildup of sucrose concentration between sieve plates and thus partially contribute to increasing the osmotic potential locally (i.e. functioning as a relay).

As conjectured from the experiments presented here, the presence of sieve plates can add the necessary structural rigidity to improve hydraulic transport efficiency within the confines of the PFH. This conjecture is explored indirectly here through idealized experiments that seek to quantify the consequences of tube expansion on sucrose front speeds. Comparing sugar front speeds measured in an ideal elastic membrane with a theoretical rigid membrane (using mathematical modeling) offers a new perspective about the controversial role of sieve plates. It will be argued here that these sieve plates act as structural dampers thereby increasing rigidity and concomitant sucrose front speed. More broadly, the experiments also offer benchmark data on tube expansion and front speeds in osmotically driven laminar flows.

## 2 Materials and methods

### 2.1 Experimental set-up

The setup here allows the quantification of osmotically driven laminar flow within closed elastic semi-permeable membrane tubes. Different membranes (representing phloem sieve tube walls) and dextrans (representing sucrose composition) were employed to cover a broad range of loading conditions on front speeds when the walls are elastic.

#### 2.1.1 Sugar

Two dextrans (Alfa Aesar, Tewksbury, MA, USA) with different molecular weights (*M_w_*) were used. Dextran J61216 and J62775 had average *M_w_* of 20 kDa (referred to as D20) and 6 kDa (referred to as D6), respectively. The initial concentration of these dextrans *c*_0_ were varied from 2 to 20 grams in 21 ml of distilled water to represent low to high osmotic potentials. These solutions were mixed with a blue dye Trypan Blue (thermoFisher Scientific, Tewksbury, MA, USA) with a *M_w_* of 960.79 Da. Even though the dye’s *M_w_* was smaller than the molecular weight cut off (MWCO) of the membranes, preliminary experiments using dyed solutions showed that the mixtures did not to leak through the membrane.

#### 2.1.2 Membrane

Two membranes with different MWCO of 3.5 kDa and 10 kDa were used with each dextran. Both membranes were semi-permeable dialysis tubes (SnakeSkin Dialysis Tubing, Thermofisher, Waltham, MA, USA) with wet initial radius of *h* = 8 mm and thickness of 0.1 mm. Before any experiment, the membrane was soaked in distilled water for an hour to limit any possibility of damaging the membrane upon stretching during the setup.

#### 2.1.3 Setup

The setup consisted of a 30 cm glass tube with rubber stopper attached at the bottom. The membrane was attached to a stopcock fitted within the rubber stopper using Parafilm and to a brass cylinder attached to a pressure transducer (PX409, Omega Engineering, Norwalk, CT, USA) at the top. Once the membrane was connected and sealed at both ends, it was inserted in the glass tube and filled with distilled water while keeping the top open to remove any pressure buildup within the membrane. At the same time, the glass tube was filled with water to keep the membrane fully hydrated. The initial length *L* of the membrane was 20.5 cm across all the experiments. Next, the dextran solution was slowly injected at the bottom with a syringe while opening the top to keep a zero pressure within the membrane as before. Once the solution was filled, the top and bottom were closed and the water in the glass tube outside the membrane was kept at constant height by attaching it to an external water reservoir (using a conical flask). A schematic figure of the setup is shown in supplementary figure S1. The movement of the sugar solution was recorded using a digital camera (Sauledo 4K, 60FPS, 48MP video recorder) acquiring images every 30 s. To ensure a reference color for later image processing, a white background was set using a white paper attached to the glass. Pressure data from the transducer was collected every 30 seconds as well.

#### 2.1.4 Data processing

Image processing tools in MATLAB programming language (Mathworks, Natick, MA) were used to detect simultaneously the front location and membrane diameter for the experiments. These data were post processed by removing the initial period (1-2 hours) that was noisy because of the inertial effects at the start of the experiment (dextran transient mixing before osmosis building up). In addition, the final time that was used to obtain the results was chosen to be the time when the pressure reached 90 % of its equilibrium value, or when the front reached the top of the membrane in case of high initial concentration. This period represented mainly the elastic regime and the initial part of the plastic regime before any membrane failure occurred after prolonged stress exposure. All in all, the study yielded 28 runs when combining both dextrans.

### 2.2 Data analysis

Two independent approaches to analyze the data were developed and applied to all 28 experiments. These approaches tackle the problem from two viewpoints that utilize differing data sets and assumptions. One approach utilizes front speeds using conservation of volume and the other is based on pressure analysis using the relation between osmotic potential and the flow of water into the membrane. The first approach is primarily kinematic and shows how the front speed is affected by allowing the membrane to expand. The second approach discusses the underlying dynamics that decrease transport efficiency due to membrane expansion.

#### 2.2.1 Conservation of volume approach

The conservation of volume can be applied on a time-dependent volume confined by the dextran concentration from the bottom spanning up to the moving front location axially and along the membrane diameter radially. The conservation of mass requires that to maintain a constant density, any increase in volume is balanced by the difference in fluxes into and out of the control volume expressed as

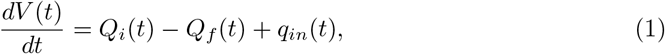

where *V* is the evolving volume of the solution within the membrane at time *t*, *Q_i_* and *Q_f_* are the axial volumetric flow rate at the entrance and exit of the control volume and *q_in_* is the radial volumetric flow rate into the volume. This radial inflow of water occurs at a dynamically changing surface area *S*(*t*) and is generated because of osmosis. It can be written as

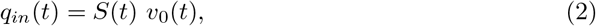

where *S*(*t*) is the surface area covered by the sugar solution and *v*_0_(*t*) is the osmotic velocity that describes the velocity of water passing radially through the membrane. The axial volumetric flow rates at inlet and front location can be approximated by their advective component and written as

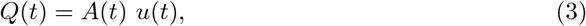

where *A*(*t*) is the cross-sectional area orthogonal to *u*(*t*) and *u*(*t*) is the axial velocity. This conservation law is analyzed over the whole period of the experiment by integrating equation (1) from initial time *t_i_* (set after removing transient effects of introducing the dextran) to final time *t_f_* (discussed earlier). In this case, the total increase of volume due to the traveling front and expanding domain can be calculated without using any pressure measurements. Assuming that the axial velocity *u_i_* at the inlet position is negligible results in

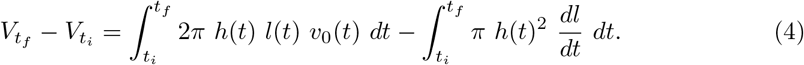

where *V_t_f__* and *V_t_i__* are the final and initial imaged volumes. To approximate the integrals on the right side of equation (4), exponential curves were fitted to the data (the change in membrane radius in time *h*(*t*) and front location *l*(*t*) ≤ *L*) using nonlinear regression. This approximation for the integrals appears consistent for all experiments except for the ones with higher dextran concentration where only a linear region was reached because the front reached the end of the tube before the development of an exponential profile. For all the experiments, both variables (i.e. *y*(*t*) = *h*(*t*), *l*(*t*)) were approximated using

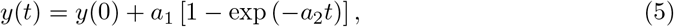

where *a*_1_ and *a*_2_ are assumed to be constant and can be determined separately from imaged *l*(*t*) and *h*(*t*) with variations in t using the nonlinear regression toolbox in MATLAB. Hence, the total radial inflow of water due to osmosis is now given by

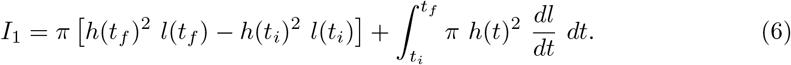

For a rigid membrane, equation (4) may be simplified to

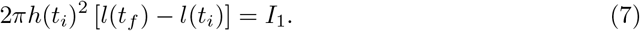

To use this approximation, the osmotic velocity *v_0_* was assumed to be equal to an effective velocity that is time independent 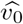 leading to

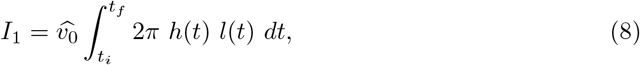

where the integral was approximated using the exponential shape for *h*(*t*) and *l*(*t*) as before. The *I*_1_ was plotted against this approximated integral (shown in supplementary figure S2A) to find the final time *t_f_* that leads to a constant osmotic velocity that can be used in both elastic and rigid membrane (shown in supplementary figure S2B). This linear period shows the range where the osmotic forcing is still approximately not far from its maximum value before the dilution effect occurs. This approximation must be viewed as a lower limit for 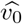 in a rigid membrane case.

#### 2.2.2 Volume-Pressure relation

The conservation law can also be applied as a point equation to relate the radius of the membrane to the total pressure within the membrane though the fluid velocity that crosses the membrane (considered here as porous media). At the membrane, the speed of the evolution of *h*(*x, t*) can be related to the fluid velocity *v_f_*(*x,t*) by

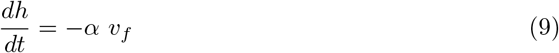

where *α* is related to the membrane elasticity and the negative sign denotes an opposite direction for the flow of water. The fluid velocity *v*_f_ is linearly proportional to the total pressure difference, which is the sum of hydrostatic and osmotic pressures, between the inside and outside of the membrane. This difference can be expressed using Darcy’s law and is given by (Darcy 1856)

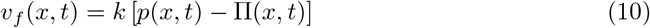

where Π is the osmotic potential, *p* is the hydrostatic pressure and *k* is the membrane permeability assumed here to be uniform in time and homogeneous in space. The osmotic potential can be further approximated using the Van’t Hoff linear relation (reasonable for low concentration)

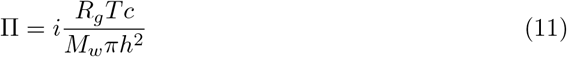

where *R_g_*, *T*, *M_w_*, *i* and *c* are the ideal gas constant, temperature (K), molecular weight of dextran (g mol^-1^), Van’t Hoff factor (that need not be the same for the two dextrans), and dextran concentration (g cm^-1^) respectively. Equation (9) can be averaged over the whole membrane to relate the average radius 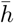 to the average hydrostatic pressure 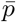 measured by the sensor (not used in the prior volume conservation analysis) and the average osmotic potential injected in the elastic membrane. When invoking Darcy’s law, this leads to (dropping the superscript for notational simplicity)

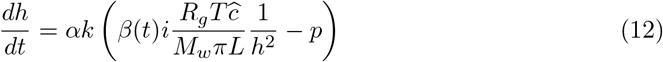

where 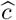 is the total amount of dextran injected in the domain in grams. The *β*(*t*) is a cutoff constant that is related to the membrane properties and later discussed. As shown in section 2.1.4, the *dh*(*t*)/*dt* is nearly uniform in space to within the camera resolution.

To understand the effect of membrane elasticity on mass flow, equation (12) was applied at the beginning and end of the experiment. Hereafter, we set the beginning of the experiment at time origin *t* = 0, and we assume that (1) the average pressure in the dialysis tube was zero (as reference) and (2) the membrane radius was the wet radius of 0.8 cm. Even though the first 1-2 hours were removed, the experiment still provided ample number of data points for statistically determining the slope of the radius profile (i.e. the speed *dh/dt*) in the linear regime. In this case, equation (12) can be written as

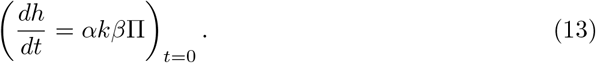

The analysis of the equation at final time (or more precisely when *t* → ∞) eliminates the rate of expansion *αk*, which is the combination of Young modulus of elasticity and membrane permeability and is shown to be similar for both membranes later on. In this case, the focus is on the cutoff coefficient at final time, which highlights the loss of efficiency in an elastic tube, and the equation is written as

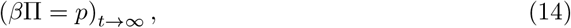

where the osmotic pressure is evaluated using the radius at equilibrium. Theoretically, if the membrane is allowed to expand without any limitation (i.e. elastic regime is maintained up to the osmotic pressure), *β* should be equal to 1 and the measured pressure should reach the osmotic potential (after including any small dilution effect). To estimate the parameters in equation (14), an exponential curve was fitted to the measured radius of the membrane and the pressure as discussed in section 2.2.1.

## 3 Results

### 3.1 Osmotic potential is less efficient in driving flow in an elastic than rigid tube

The volume-pressure relation shows a decrease of efficiency of the osmotic potential when the cutoff coefficient *β* is evaluated at initial and final time of the experiment. At maximum efficiency, *β* should be unity. However, this was not the case for the described experiments. Figure 1A shows the initial rate of expansion of the elastic membrane plotted against the initial osmotic pressure. The inset plot shows the same results when the Van’t Hoff factor *i* was assumed to be unity for both dextrans. Assuming that both membranes behaved in a similar manner with each dextran because of similar MWCO to molecular weight ratio, the different slope between both dextrans can only be attributed to the Van’t hoff factor (as in the inset). Figure 1A repeats the inset result when i was corrected for D20. The revised relation was approximately linear meaning that the membrane property *αk* and the initial cutoff *β*(0) were constant as expected. The supplemental figure S3 also shows this result when (*dh/dt* Π)_*t*_=0 was plotted against Π(0) leading to approximately a constant value. Figure 1B shows the average pressure within the membrane plotted against the osmotic pressure both at equilibrium (but using the corrected Van’t Hoff factor). The cutoff coefficient *β*(*t* → ∞) was not equal to one since the pressure never attained its theoretical value even when dilution effects were included. It is later shown that the difference in i here is also compatible with the conservation of volume approach.

**Figure 1:**
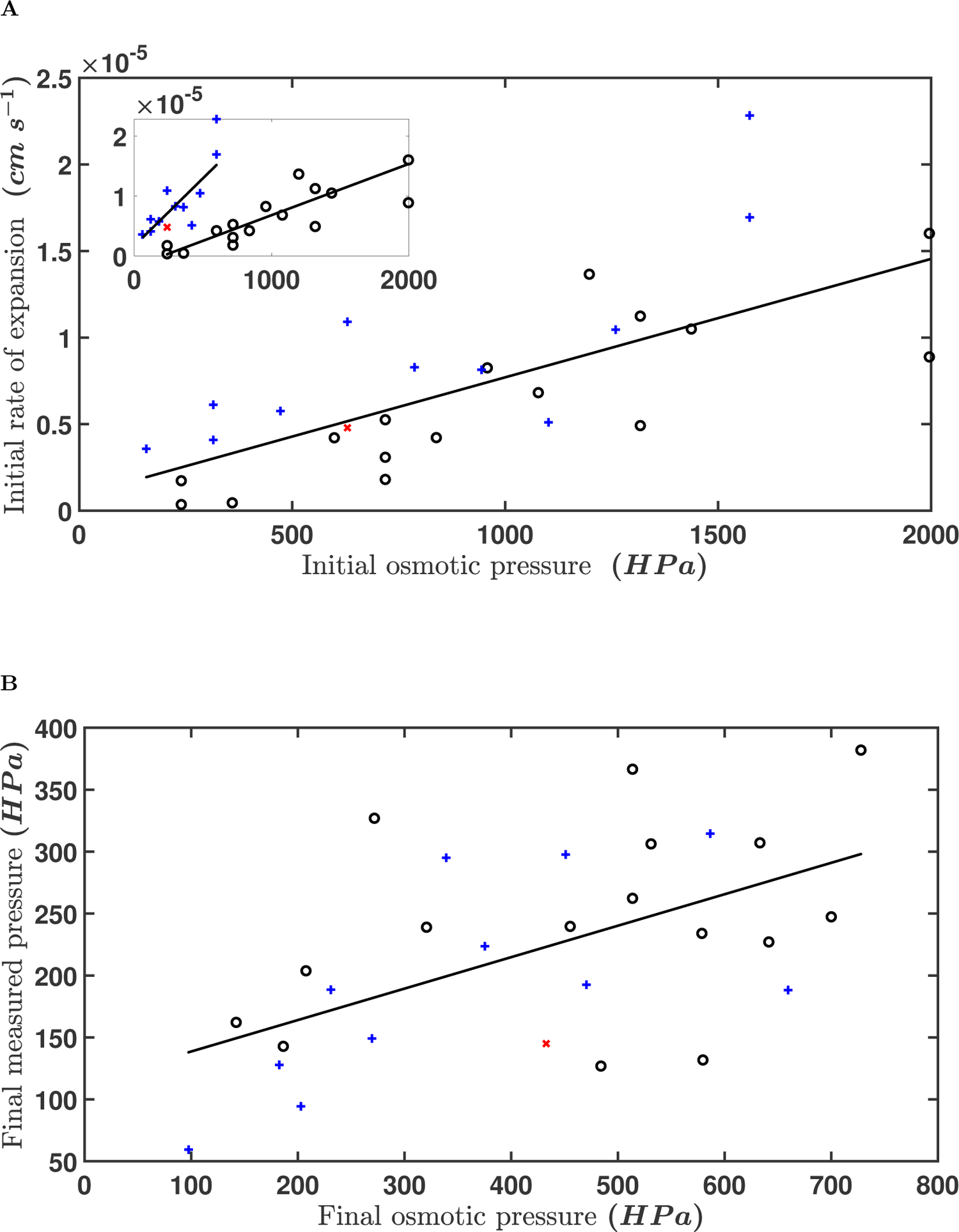
(*a*) Initial rate of expansion (*dh/dt* at *t* = 0) as a function of the initial osmotic pressure Π_*i*_ using the corrected Van’t Hoff coefficient *i* (coefficient of determination *r*^2^ = 0.54 and regression slope *p* = 0.65). Inset plot shows the results using a Van’t Hoff coefficient set to unity. (*b*) Measured pressure as a function of the calculated osmotic pressure at final time *t_f_* for both dextrans (*r*^2^ = 0.3 and *p* = 0.004). Black circles denote D6, blue crosses denote D20 and red star denote D20 with a 3.5 K membrane.

Figure 2 shows the results for the cutoff coefficient at final time while including the corrected *i* (calculated from figure 1A). In figure 2A, *β* was plotted against the osmotic pressure at final time, which includes the dilution effects of the expanded membrane. As expected, the cutoff coefficient is inversely dependent on the osmotic potential. It also shows that at higher osmotic potential (when the maximum deformation of the membrane was reached), this coefficient showed a decrease of efficiency between 0.4 and 0.6 of the maximum osmotic potential. In figure 2B, *β* was plotted against the initial expansion rate (*dh/dt* at *t_i_*) of the membrane. This shows how *β* depends on the maximum work done against the membrane, which occurred at *t_i_* because of the maximum osmotic potential. Yet again, a decrease of efficiency is seen because some work derived from the osmotic pressure was spent on membrane expansion. The coefficient varied between 0.4 and 0.6 for most of the experiments supporting findings of section 3.2 shown next.

**Figure 2:**
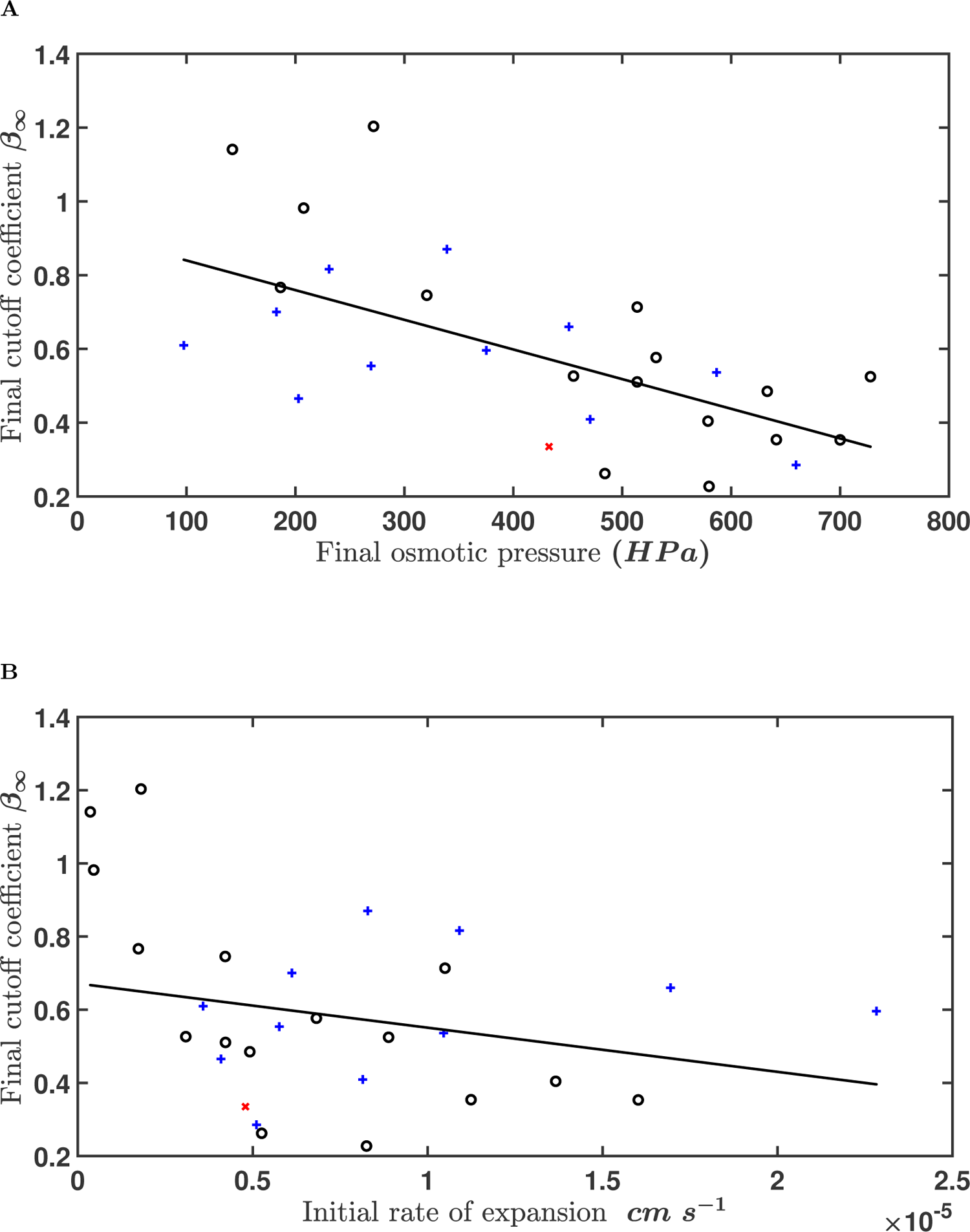
Cutoff coefficient *β* at final time *t_f_* as a function of (a) the osmotic pressure Π_*i*_ at final time *t_f_* (*r*^2^ = 0.42 and *p* = 0.0009) and (b) the initial rate of expansion (*dh/dt* at *t* = 0) for both dextrans using the corrected Van’t Hoff coefficient (*r*^2^ = 0.09 and *p* = 0.21). Black circles denote D6, blue crosses denote D20 and red star denote D20 with a 3.5 K membrane.

### 3.2 Sugar front travels faster in a rigid tube

The conservation of volume analysis shows that the Reynolds number (that linearly depends on the osmotic velocity) increases as a function of the initial osmotic pressure and the increase was faster in the case of crosses than for circles (figure 3A). The flow of water through the membrane is here presented in non-dimensional form using a Reynolds number 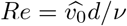, where *d* is the thickness of the membrane representing the travel distance across the membrane and *ν* is the kinematic viscosity of the background water being pulled into the membrane (distilled). It was plotted against the average osmotic potential, which is the driving force for the flow, using the initial radius of the membrane at rest. As in section 3.1, the difference in the slopes was attributed to the fact that *i* ≠ 1 for D20. The ratio between both slopes resulted in a Van’t hoff factor of ≈ 2.6, which was similar to the result found in section 3.1 (≈ 2.8). This is because both membranes behave similarly as shown in figure 3B. This difference in *i* was corrected for D20 and both are plotted in figure 3A. This figure clearly shows that the assumption to use a linear Darcy law and Van’t hoff approximations are valid for this range of sugar concentrations since the flow into the membrane is slow (Re ≪ 1), and the speed increases linearly with the driving force.

**Figure 3:**
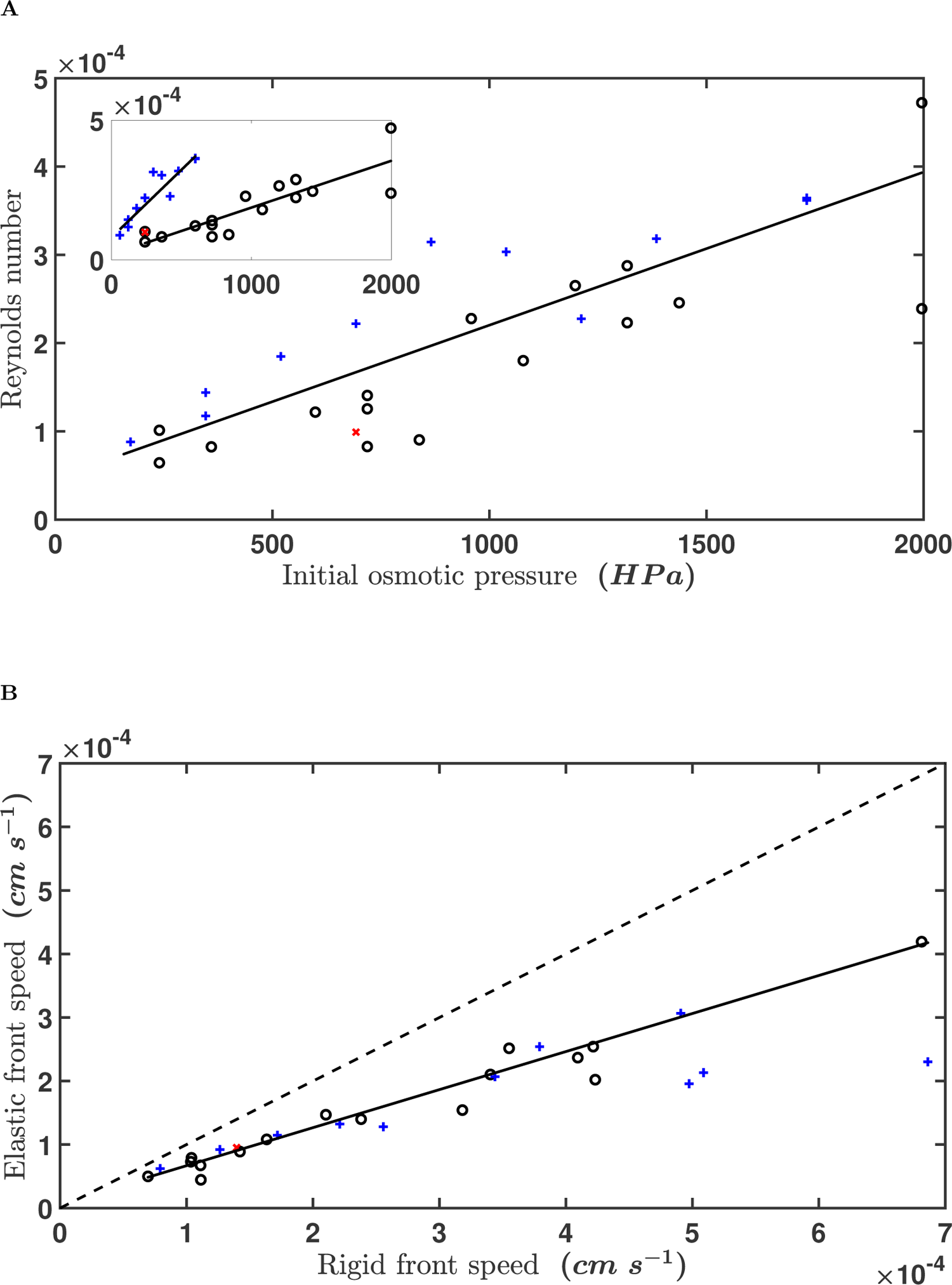
(*a*) Reynolds number (*Re* = *v*_0_*d*/*ν*) as a function of the initial osmotic pressure Π_*i*_ using the corrected Van’t Hoff coefficient *i*. Inset plot shows the results using a Van’t Hoff coefficient set to unity (*r*^2^ = 0.64 and *p* = 2 × 10^-7^). (*b*) Measured front speed for an elastic membrane as a function of the calculated of an hypothetical rigid membrane at final time *t_f_* for both dextrans (*r*^2^ = 0.69 and *p* = 4 × 10^-20^). Solid dashed line shows the 1×1 plot. Black circles denote D6, blue crosses denote D20 and red star denote D20 with a 3.5 K membrane.

The findings of the conservation of volume analysis shows that around 40% of the osmotic potential was lost due to membrane expansion where some of the work was exerted on moving the membrane uniformly in the radial direction instead of generating excess pressure to move the dextran front in the axial direction. As shown in figure 3B, both dextrans with their respective membranes behaved similarly where there is a loss of efficiency compared to a rigid tube. The dashed line shows the one-to-one curve and the data are lying on the fitted curve with a slope of ≈ 0.59. This slope agrees with the independent result of section 3.1. This figure also suggested that the attribution of the different slopes in the inset of figure 3A to *i* is reasonable since they behaved similarly from a conservation perspective where all data points converged on the same fitted line. In this figure, the measured sugar front speed for an elastic tube was plotted against the calculated sugar front speed for an assumed rigid tube. These front speeds were calculated over the whole period of the experiment. The front location at final time for the rigid tube was calculated using equation (7) and was measured from data for the elastic tube. The linearity in figure 3A also allows the use of the same average osmotic velocity 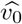 for both cases shown in figure 3B.

## 4 Discussion

The efficiency of solute transport in osmotically driven flow within a semi-permeable elastic membrane was studied using experiments for different osmotic potential. Using two different approaches, the results show that membrane rigidity increases the efficiency of solute transport in osmotically driven flow. In the first approach, front speeds extracted from the experiments were compared to front speeds of a hypothetical rigid membrane calculated using conservation of volume. This approach showed that part of the osmotic potential is lost due to the work done on the membrane. In the second approach, pressure-volume (or radius) relation showed that, in addition to the loss of energy due to the work done to expand the membrane, the efficiency of solute transport decreased because of dilution effect. For osmotically driven flow, the driving force (i.e. the osmotic potential) depends linearly on the average concentration within the membrane through the Van’t Hoff relation. This average concentration depends inversely on the cross-sectional area of the membrane (i.e. Π ~ *h*^-2^). When the membrane is allowed to expand, the cross-sectional area increases leading to a decrease in the osmotic potential available to drive the flow. To the contrary, a rigid membrane whose cross-sectional area does not change and sustain the same osmotic potential over the whole time will experience faster front speeds. These results can be used to provide new conjectures on the role of sieve plates inside the phloem. Previous theories have attributed the role of sieve plates be strictly hydraulic. It was argued that increase in efficiency may be achieved through decreasing the distance over which sugar is being transported, also called the relay effect (Lang 1979). Here, we hypothesize that sieve plates can act as structural dampers. The added rigidity translates to hydraulic efficiency in return by allowing excess osmotic potential to contribute to hydraulic head gradient moving water and sugars but with minor hydraulic headloss.

## Funding

This work was supported by the U.S. National Science Foundation [NSF-IOS-1754893, NSF-AGS-2028633], the Department of Energy [DE-SC0022072] and Los Alamos Directed Research and Development Exploratory Research Grant [No. 2020109DR].

## Data accessibility

Data is available on Duke Research Data Repository (https://doi.org/10.7924/r45d8zp3s).

## Notes

https://doi.org/10.7924/r45d8zp3s

### Competing Interest Statement

The authors have declared no competing interest.

### Summary of Updates

Data has been added

## References

Cabrita, P., Thorpe, M. & Huber, G. (2013), ‘Hydrodynamics of steady state phloem transport with radial leakage of solute’, Frontiers in Plant Science 4, 531.

Darcy, H. (1856), Les Fontaines publiques de la ville de Dijon. Exposition et application des principes à suivre et des formules à employer dans les questions de distribution d’eau, etc, V. Dalamont.

Jensen, K. (2018), ‘Phloem physics: mechanisms, constraints, and perspectives’, Current Opinion in Plant Biology 43, 96–100.

Jensen, K., Berg-Sørensen, K., Friis, S. M. & Bohr, T. (2012), ‘Analytic solutions and universal properties of sugar loading models in Münch phloem flow’, Journal of Theoretical Biology 304, 286–296.

Jensen, K., Rio, E., Hansen, R., Clanet, C. & Bohr, T. (2009), ‘Osmotically driven pipe flows and their relation to sugar transport in plants’, Journal of Fluid Mechanics 636, 371–396.

Kalmbach, L. & Helariutta, Y. (2019), ‘Sieve plate pores in the phloem and the unknowns of their formation’, Plants 8(2), 25.

Lang, A. (1979), ‘A relay mechanism for phloem translocation’, Annals of Botany 44(2), 141–145.

Lucas, W. J., Ding, B. & van der Schoot, C. (1993), ‘Tansley review no. 58. plas-modesmata and the supracellular nature of plants’, New Phytologist pp. 435–476.

Münch, E. (1930), Stoffbewegungen in der Pflanze, G. Fischer, Jena, Germany.

Nakad, M., Domec, J.-C., Sevanto, S. & Katul, G. (2022), ‘Radial–axial transport coordination enhances sugar translocation in the phloem vasculature of plants’, Plant Physiology

Nikinmaa, E., Hölttä, T., Hari, P., Kolari, P., Mäkelä, A., Sevanto, S. & Vesala, T. (2013), ‘Assimilate transport in phloem sets conditions for leaf gas exchange’, Plant, Cell & Environment 36(3), 655–669.

Phillips, R. & Dungan, S. (1993), ‘Asymptotic analysis of flow in sieve tubes with semi-permeable walls’, Journal of Theoretical Biology 162(4), 465–485.

Savage, J., Beecher, S., Clerx, L., Gersony, J., Knoblauch, J., Losada, J., Jensen, K., Knoblauch, M. & Holbrook, N. (2017), ‘Maintenance of carbohydrate transport in tall trees’, Nature Plants 3(12), 965.

Sevanto, S. (2014), ‘Phloem transport and drought’, Journal of experimental botany 65(7), 1751–1759.

Sevanto, S., Mcdowell, N. G., Dickman, L. T., Pangle, R. & Pockman, W. T. (2014), ‘How do trees die? a test of the hydraulic failure and carbon starvation hypothe-ses’, Plant, cell & environment 37(1), 153–161.

Sevanto, S., Ryan, M., Dickman, L. T., Derome, D., Patera, A., Defraeye, T., Pan-gle, R. E., Hudson, P. J. & Pockman, W. T. (2018), ‘Is desiccation tolerance and avoidance reflected in xylem and phloem anatomy of two coexisting arid-zone coniferous trees?’, Plant, Cell & environment 41(7), 1551–1564.

Tateshima, S., Chien, A., Sayre, J., Cebral, J. & Viñuela, F. (2010), ‘The effect of aneurysm geometry on the intra-aneurysmal flow condition’, Neuroradiology 52(12), 1135–1141.

Thompson, M. & Holbrook, N. (2003), ‘Application of a single-solute non-steady-state phloem model to the study of long-distance assimilate transport’, Journal of Theoretical Biology 220(4), 419–455.

