## Supplemental figures for "Experiments on osmotically driven flow in idealized elastic membranes"

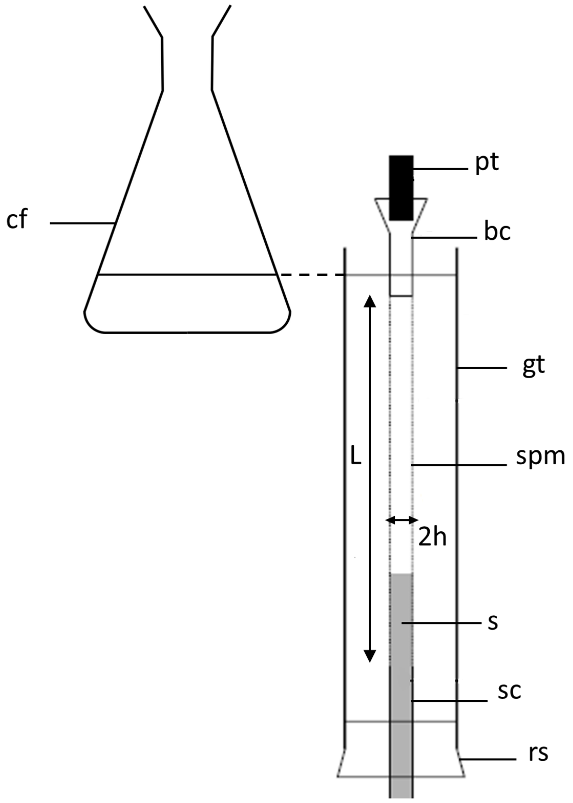


Figure S1: Experimental setup used to track the front speed and volume expansion. (s) sugar solution, (sc) stopcock, (rs) rubber stopper, (spm) semi-permeable membrane, (gt) glass tube, (bc) brass cylinder, (pt) pressure transducer and (cf) conical flask. The $L = 20.5$ cm and $h$ are the length and radius of the membrane respectively.

**A**

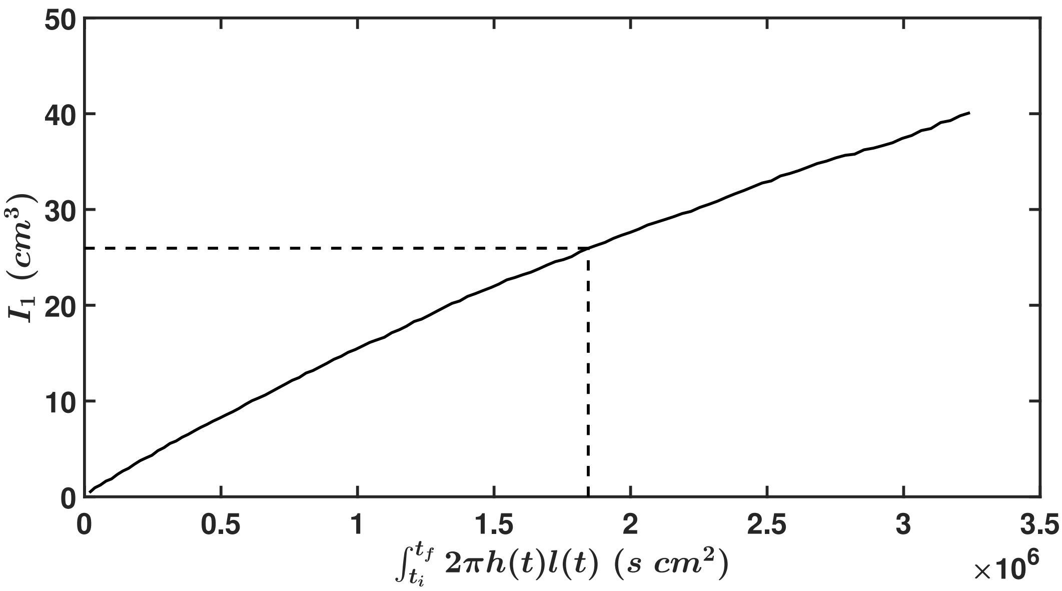


**B**

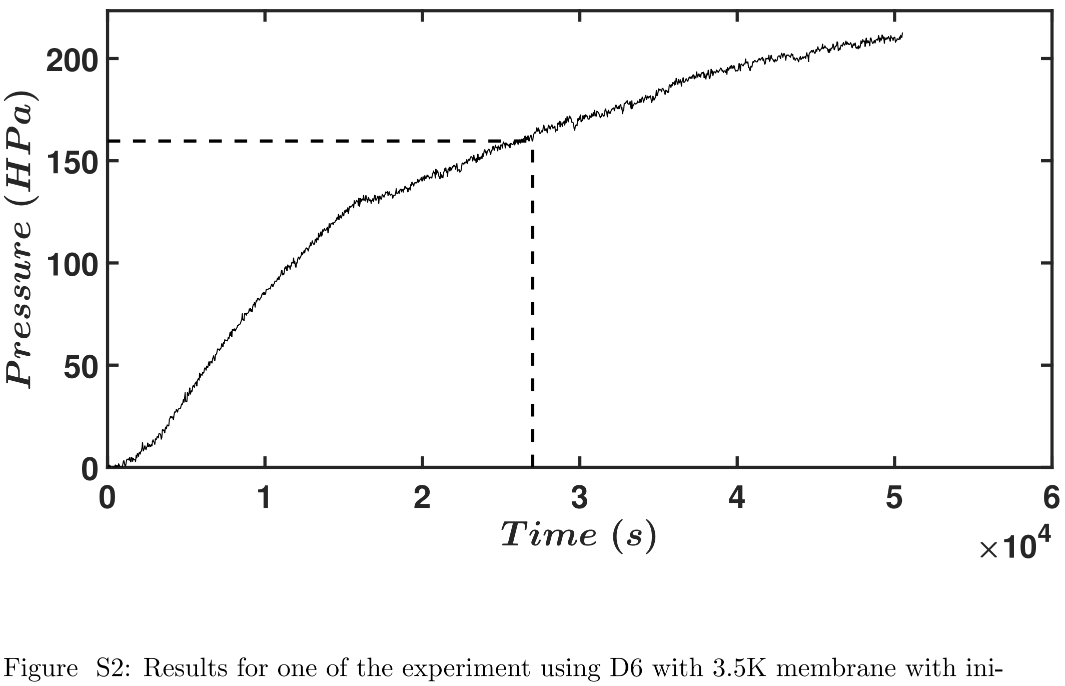


Figure S2: Results for one of the experiment using D6 with 3.5K membrane with initial concentration equal to 7.2 g. ($a$) The total inflow of water due to osmosis $I_1$ plotted against the approximated integral $\int_{t_i}^{t_f} 2\pi\ h(t)\ l(t)\ dt$. ($b$) Measured pressure plotted against time. Dashed lines denote the period where a constant osmotic velocity $v_0$ is still valid.

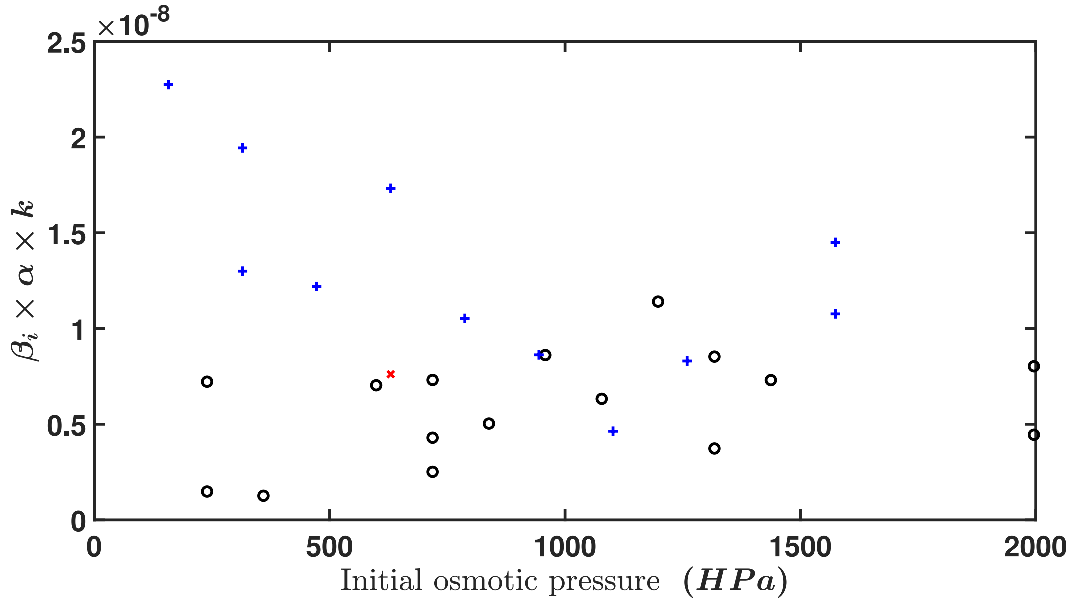


Figure S3: Calculated coefficient $\beta\alpha k$ as a function of the calculated osmotic pressure at intial time for both dextrans. Black circles denote D6, blue crosses denote D20 and red star denote D20 with a 3.5 K membrane.
